# Leveraging shared connectivity to aggregate heterogeneous datasets into a common response space

**DOI:** 10.1101/741975

**Authors:** Samuel A. Nastase, Yun-Fei Liu, Hanna Hillman, Kenneth A. Norman, Uri Hasson

## Abstract

Connectivity hyperalignment can be used to estimate a single shared response space across disjoint datasets. We develop a connectivity-based shared response model that factorizes aggregated fMRI datasets into a single reduced-dimension shared connectivity space and subject-specific topographic transformations. These transformations resolve idiosyncratic functional topographies and can be used to project response time series into shared space. We evaluate this algorithm on a large collection of heterogeneous, naturalistic fMRI datasets acquired while subjects listened to spoken stories. Projecting subject data into shared space dramatically improves between-subject story time-segment classification and increases the dimensionality of shared information across subjects. This improvement generalizes to subjects and stories excluded when estimating the shared space. We demonstrate that estimating a simple semantic encoding model in shared space improves between-subject forward encoding and inverted encoding model performance. The shared space estimated across all datasets is distinct from the shared space derived from any particular constituent dataset; the algorithm leverages shared connectivity to yield a consensus shared space conjoining diverse story stimuli.

**Highlights:**

- Connectivity SRM estimates a single shared space across subjects and stimuli
- Topographic transformations resolve idiosyncrasies across individuals
- Shared connectivity space enhances spatiotemporal intersubject correlations
- Semantic model-based encoding and decoding improves across subjects
- Transformations project into a consensus space conjoining diverse stimuli

## Introduction

The developing infrastructure for data sharing (alongside evolving incentives) has led to a proliferation of publicly available “open” neuroimaging data. Although still overshadowed by traditional task and resting-state acquisitions, we are beginning to see more public data collected during rich, naturalistic paradigms (DuPre et al., 2019; Hanke et al., 2016, e.g., 2014; Taylor et al., 2017). Although the neuroimaging community unequivocally benefits from the increasing availability of public data (Milham et al., 2018; Poldrack and Gorgolewski, 2014), this trend introduces a challenge. Namely, datasets are often markedly heterogeneous—i.e., collected on different scanners, using different acquisition parameters, with different samples of subjects—and require sophisticated harmonization (e.g., Yamashita et al., 2019). Furthermore, in the context of naturalistic stimuli (e.g., movie-watching, story-listening), stimuli vary considerably from experiment to experiment. Here we focus on a particular aspect of harmonization: finding a shared functional response space across heterogeneous naturalistic datasets.

In order to fully realize the potential of “big” neuroimaging data for prediction and translational purposes, we need to obtain some level of correspondence across individuals (Dubois and Adolphs, 2016; Gabrieli et al., 2015; Woo et al., 2017). Typically, each individual brain is spatially normalized to a standard space based on macroanatomical features such as sulcal curvature (Coalson et al., 2018; Fischl et al., 1999). However, fine-grained functional response topographies (e.g., Brett et al., 2002; Duncan et al., 2009; Frost and Goebel, 2012; Haxby et al., 2014; Zhen et al., 2017, 2015) and connectivity patterns (e.g., Bijsterbosch et al., 2019; Braga and Buckner, 2017; Gordon et al., 2017; Langs et al., 2016) are not tightly coupled to macroanatomical features and are markedly idiosyncratic across individuals. Information encoded at this finer scale may be inaccessible based on anatomical alignment alone (Feilong et al., 2018; Kong et al., 2019). In order to leverage large volumes of data for prediction in individuals, we need to resolve these idiosyncrasies in functional–anatomical correspondence. Hyperalignment is a family of algorithms for normalizing functional data into a common space by resolving topographic idiosyncrasies (Guntupalli et al., 2016; Haxby et al., 2011). These methods hinge on functional commonalities to drive normalization—typically a rich stimulus is used to evoke stereotyped, time-locked response trajectories across subjects. However, recent work by Guntupalli and colleagues (2018) demonstrates that functional connectivity can be used to effectively drive functional normalization absent any shared stimulus. Each voxel’s participation in functional networks elsewhere in the brain can provide a shared functional signature sufficient for resolving topographic idiosyncrasies. One important product of this development, which the authors examined in detail, is the extension of hyperalignment to resting-state functional connectivity. In this case, the “rest” task yields rich and consistent enough connectivity patterns to support functional normalization. A separate, largely unexplored avenue is that connectivity hyperalignment may allow us to define a single common space across distinct naturalistic stimuli or tasks.

Here, we use a variant of connectivity-based hyperalignment (Guntupalli et al., 2018) to showcase the utility of aggregating disjoint naturalistic story-listening datasets into a single, shared response space. For a given region of interest (ROI), we first compute intersubject functional correlations (ISFC; Simony et al., 2016) between each voxel and a set of parcels tiling the cortex (i.e., connectivity targets; Glasser et al., 2016). We then apply the shared response model (SRM; Chen et al., 2015) to these connectivity patterns to find a reduced-dimension connectivity space shared across both subjects and stimuli. Critically, in the context of a task such as listening to spoken stories, we expect these coarse connectivity patterns to be well-preserved across subjects and stimuli. The SRM effectively decomposes the connectivity data across all datasets into a shared connectivity space, and a set of subject-specific transformation matrices that resolve topographic idiosyncrasies. Although the shared model is derived from functional connectivity, the subject-specific topographic transformations can be used to project response time series into shared space. We benchmark this algorithm on a large, heterogeneous collection of story-listening functional MRI datasets assembled over the course of approximately seven years. This data collection comprises 10 unique auditory story stimuli across 300 scans with 160 unique subjects. We evaluate the shared space using between-subject time-segment classification (e.g., Haxby et al., 2011), temporal and spatial intersubject correlations (e.g., Nastase et al., 2019), and between-subject semantic model-based encoding and decoding (e.g., Huth et al., 2016).

## Materials and methods

### Participants

We aggregated fMRI datasets collected between 2011 and 2018 comprising 10 story stimuli and 160 subjects totalling 300 scans (mean age = 22.5 years, *SD* = 6.25, range: 18–53; 89 reported female). Subjects with behavioral comprehension scores (where applicable) lower than 25% accuracy were excluded. Furthermore, we computed leave-one-subject-out ISCs in a left early auditory cortex ROI (Glasser et al., 2016) for all subjects in each dataset at temporal lags ranging from –100 to 100 TRs and excluded any subjects with a peak ISC at lags exceeding ±1 TR. Here we briefly summarize the resulting sample size and demographics for each dataset, and point to previously published work using these data (see Table 1). The datasets are named according to the names of the corresponding story stimuli, with abbreviated aliases used in analysis and figures. The “Pie Man” data (alias: *pieman*) comprised 46 subjects (mean age = 22.4 years, *SD* = 3.8, 23 reported female; (Simony et al., 2016). The “Pretty Mouth and Green My Eyes” data (alias: *prettymouth*) comprised 19 subjects (mean age = 20.2 years, SD = 2.1, 9 reported female) from the “cheating” condition of the context manipulation described by Yeshurun and colleagues (2017b). The “Milky Way” data comprised 16 subjects (mean age = 20.0 years, *SD* = 1.5, 7 reported female) from one condition (Story1) of the word-substitution manipulation described by Yeshurun and colleagues (2017a). The previously unpublished “Slumlord” and “Reach for the Stars One Small Step at a Time” stories were presented in a single scanning run (alias: *slumlordreach*) and comprised 16 subjects (mean age = 21.13 years, *SD* = 2.4, 8 reported female). The previously unpublished “It’s Not the Fall That Gets You” data (alias: *notthefall*) comprised 18 subjects (mean age = 21.4 years, *SD* = 2.5, 8 reported female). The previously unpublished “The 21st Year” data (alias: *21styear*) comprised 26 subjects (mean age = 23.3 years, SD = 6.7, 14 reported female). Finally, two stories recorded at the Princeton Neuroscience Institute (PNI) served as stimuli: “Pie Man (PNI)” (alias: *pieman (PNI)*) and “Running from the Bronx (PNI)” (alias: *bronx (PNI)*). The “Pie Man (PNI)” data comprised 39 subjects (mean age = 23.3 years, *SD* = 7.7, 28 reported female). The “Running from the Bronx (PNI)”, “I Knew You Were Black” (alias: *black*), and “The Man Who Forgot Ray Bradbury” (alias: *forgot*) data were collected at the same time and comprised roughly the same sample of 40 subjects (mean age = 23.3 years, *SD* = 7.6, 29 reported female; (Lin et al., 2019).

**Table 1:**
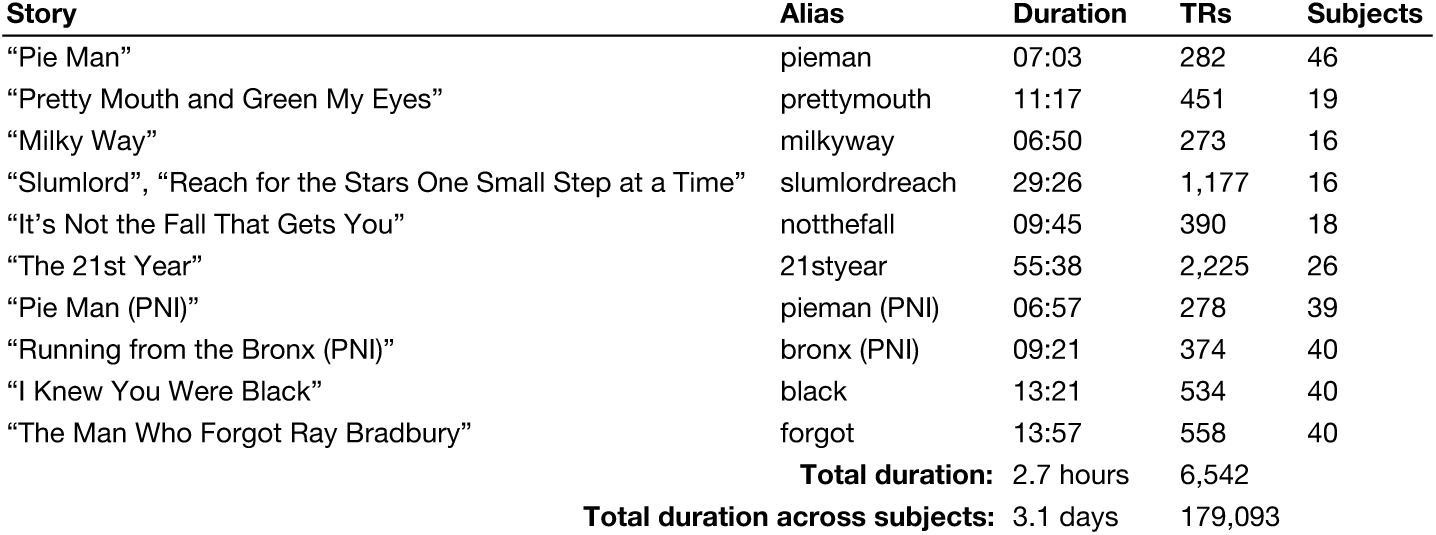
Summarization of 10 benchmark story-listening fMRI datasets.

Stimuli comprised 10 spoken stories. In addition to the names of the stories, we use abbreviated aliases in analysis and figures. Story durations are listed in “minutes:seconds” format, and exclude any silence or music bookending the story itself. The number of TRs for each story also excludes any TRs corresponding to silence or music; a 1.5-second TR was used for all acquisitions. The sample size listed for each story corresponds to the number of subjects used for subsequent analyses after applying exclusion criteria (see Participants section). The “total duration” is simply the sum of story durations; i.e., the duration of unique stimuli across datasets (not accounting for the number of subjects in each dataset). The “total duration across subjects” takes into account the number of subjects acquired for each story and reflects the grand total duration if all data were concatenated across both subjects and stories. Note that “Slumlord” and “Reach for the Stars One Small Step at a Time” are distinct stories, but were presenting one after the other in a single scanning run. The “Pie Man (PNI)” and “Running from the Bronx (PNI)” stimuli were recorded while the speaker underwent an fMRI scan, and those have relatively low audio quality. The “Pie Man (PNI)” stimulus recorded at PNI differs from the original “Pie Man” stimulus recorded at a live storytelling event.

### Stimuli and design

Story stimuli were presented auditorily and ranged from ∼7–56 minutes in duration (summarized in Table 1). The stimuli including from professional storytellers performing for a live audience, actors performing written narratives, and authors reading their written works. For each dataset, TRs corresponding to the story stimulus were isolated by discarding any TRs corresponding to silence or music (padding the beginning or end of a run). In general, participants were instructed to maintain fixation on a centrally-presented crosshair or dot and listen to the story. Behavioral questionnaires assessing narrative comprehension were acquired for the “Pretty Mouth and Green My Eyes”, “Milky Way”, “Slumlord” and “Reach for the Stars One Small Step at a Time”, “The 21st Year”, “Pie Man (PNI)”, “Running from the Bronx (PNI)”, “I Knew You Were Black”, and “The Man Who Forgot Ray Bradbury”. These comprehension scores were used to initially exclude poor-performing or noncompliant participants (participants with accuracies lower than 25%), but were not used in subsequent analyses.

### Image acquisition

MRI data for the “Pie Man”, “Pretty Mouth and Green My Eyes”, “Milky Way”, “Slumlord”, “Reach for the Stars One Small Step at a Time”, “It’s Not the Fall that Gets You”, and “The 21st Year” were collected using a 3T Siemens Skyra with a 20-channel phased-array head coil. Functional blood-oxygenation-level-dependent (BOLD) images were acquired in an interleaved fashion using gradient-echo echo-planar imaging with an in-plane acceleration factor of 2 using mSENSE: TR/TE = 1500/28 ms, flip angle = 64°, bandwidth = 1445 Hz/Px, in-plane resolution = 3 × 3 mm, slice thickness = 4 mm, matrix size = 64 × 64, FoV = 192 × 192 mm, 27 axial slices with roughly full brain coverage and no gap, anterior–posterior phase encoding. At the beginning of each run, three dummy scans were acquired and discarded by the scanner to allow for signal stabilization. T1-weighted structural images were acquired using a high-resolution single-shot MPRAGE sequence with an in-plane acceleration factor of 2 using GRAPPA: TR/TE/TI = 2300/3.08/900 ms, flip angle = 9°, bandwidth = 240 Hz/Px, in-plane resolution 0.859 × 0.859 mm, slice thickness 0.9 mm, matrix size = 256 × 256, FoV = 172.8 × 219.9 × 219.9 mm, 192 sagittal slices, ascending acquisition, no fat suppression.

MRI data for the “Pie Man (PNI)”, “Running from the Bronx (PNI)”, “I Knew You Were Black”, and “The Man Who Forgot Ray Bradbury” stores were collected using a 3T Siemens Prisma with a 64-channel head coil. Functional images were acquired in an interleaved fashion using gradient-echo echo-planar imaging with a multiband acceleration factor of 3 and no in-plane acceleration: TR/TE 1500/31 ms, flip angle = 67°, bandwidth = 2480 Hz/Px, in-plane resolution = 2.5 × 2.5 mm, slice thickness 2.5 mm, matrix size = 96 × 96, FoV = 240 × 240 mm, 48 axial slices with full brain coverage and no gap, anterior–posterior phase encoding, three dummy scans. T1-weighted structural images were acquired using a high-resolution single-shot MPRAGE sequence with an in-plane acceleration factor of 2 using GRAPPA: TR/TE/TI = 2530/3.3/1100 ms, flip angle = 7°, bandwidth = 200 Hz/Px, in-plane resolution 1.0 × 1.0 mm, slice thickness 1.0 mm, matrix size = 256 × 256, FoV = 176 × 256 × 256 mm, 176 sagittal slices, ascending acquisition, no fat suppression, X min X s total acquisition time. T2-weighted structural images were acquired using a high-resolution single-shot MPRAGE sequence with an in-plane acceleration factor of 2 using GRAPPA: TR/TE = 3200/428 ms, flip angle = 120°, bandwidth = 200 Hz/Px, in-plane resolution 1.0 × 1.0 mm, slice thickness 1.0 mm, matrix size = 256 × 256, FoV = 176 × 256 × 256 mm, 176 sagittal slices, ascending acquisition, no fat suppression.

### Preprocessing

All MRI data were preprocessed using fMRIPrep (Esteban et al., 2019), which uses Nipype (Gorgolewski et al., 2011) to adaptively construct workflows based on metadata. Anatomical T1-weighted images were corrected for intensity non-uniformity (Tustison et al., 2010) and skull-stripped based on the OASIS template using ANTs (Avants et al., 2008). Cortical surfaces were reconstructed using FreeSurfer (Dale et al., 1999) and tissue segmentation was performed using FSL (Zhang et al., 2001). T2-weighted images were also supplied to surface reconstruction where applicable.

Functional data were slice-time corrected using AFNI (Cox, 1996) and motion corrected using FSL (Jenkinson et al., 2002). “Fieldmap-less” susceptibility distortion correction was performed by co-registering the functional image to the T1-weighted image for that subject with intensity inverted (Wang et al., 2017) constrained with an average field map template (Treiber et al., 2016) using ANTs. Functional images were next co-registered to the corresponding T1-weighted image using FreeSurfer’s boundary-based registration (Greve and Fischl, 2009). Transformations for performing motion correction, susceptibility distortion correction, and functional to anatomical registration were concatenated and applied in a single step with Lanczos interpolation using ANTs. Functional data were then resampled to the subject-specific cortical surface models by averaging samples at six intervals along the normal between the white matter and pial surfaces. Functional data were then spatially normalized to the fsaverage surface template based on sulcal curvature and downsampled to the fsaverage6 template (Fischl et al., 1999). All subsequent analyses (including functional normalization) were performed on surface data (Coalson et al., 2018), and functional normalization algorithms are compared to relatively high-performing nonlinear surface-based anatomical normalization (Klein et al., 2010). Note that the terms “voxel” and “vertex” are effectively interchangeable for the analyses of interest; although we sometimes refer to voxels in keeping with conventions in the literature, all analyses of interest were performed explicitly on surface vertices.

The following confound variables were regressed out of the signal In a single step (Lindquist et al., 2019) using AFNI’s 3dTproject: linear and quadratic trends, sine/cosine bases for high-pass filtering (cutoff: 0.00714 Hz; ∼140 s), six head motion parameters and their derivatives, framewise displacement (Power et al., 2014), and six principal component time series from anatomically-defined cerebrospinal fluid and white matter segmentations (Behzadi et al., 2007).

### Regions of interest

We evaluated the shared model in several regions of interest (ROIs) defined according to a multimodal parcellation (MMP) based on anatomical and functional data from the Human Connectome Project (Glasser et al., 2016). The surface-based parcellation was projected to the fsaverage surface template and downsampled to the fsaverage6 template (Mills, 2016). We focused on four large cortical regions (Fig. 1), each of which subtends several smaller cortical areas. Following the MMP, early auditory cortex (EAC) comprised five areas (A1, MBelt, LBelt, PBelt, RI) and contained 808 and 638 vertices in the left and right hemispheres, respectively. Auditory association cortex (AAC) comprised eight areas (A4, A5, STSdp, STSda, STSvp, STSva, STGa, TA2) and contained 1,420 (left hemisphere) and 1,493 (right hemisphere) vertices. The temporo-parieto-occipital junction (TPOJ) comprised five areas (TPOJ1, TPOJ2, TPOJ3, STV, PSL) and contained 847 (left hemisphere) and 1,188 (right hemisphere) vertices. To reduce the posterior cingulate cortex region to a more comparable size (originally 14 areas containing over 2,500 vertices per hemisphere), we selected seven core areas (POS1, POS2, v23ab, d23ab, 31pv, 31pd, 7m) containing 1,198 (left hemisphere) and 1,204 (right hemisphere) vertices; we refer to this region as posterior medial cortex (PMC). These ROIs (EAC, AAC, TPOJ, and PMC) span a cortical hierarchy supporting language and narrative comprehension (Baldassano et al., 2017; Huth et al., 2016; Lerner et al., 2011). The selection of ROIs was not intended to be exhaustive; rather, we aimed to benchmark the shared model in a sample of relevant ROIs ranging from low-level sensory cortex to high-level association cortex. We analyze ROIs in both hemispheres (left and right) separately in all subsequent analyses, but generally collapse across hemispheres for statistical summarization.

**Fig. 1.**
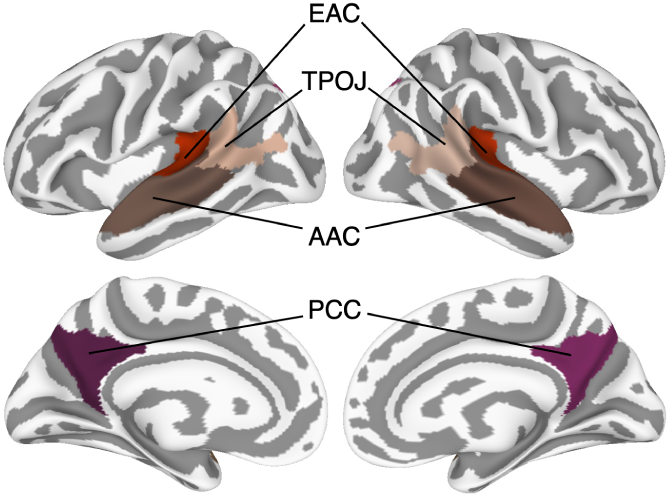
Regions of interest. Four large cortical regions roughly capturing the processing hierarchy for language and narrative comprehension were defined according to a multimodal parcellation (Glasser et al., 2016): early auditory cortex (EAC), auditory association cortex (AAC), temporo-parieto-occipital junction (TPOJ), and posterior medial cortex (PMC).

### Connectivity-based shared response model

Here we apply a variant of connectivity hyperalignment to multiple datasets with largely non-overlapping subjects, where all datasets share a common task (i.e., story-listening) and each “dataset” corresponds to a unique, naturalistic stimulus (auditory recordings of spoken stories). We use the term “hyperalignment” (via Haxby et al., 2011) to refer to the superordinate class of functional normalization algorithms that leverage commonality of function to transform subject-specific, topographic responses into a common response space. The current work develops a specific algorithm within this overarching class, which we refer to as connectivity-based shared response model (connectivity SRM or cSRM; Fig. 2). Note, however, that this algorithm differs in several ways from the core implementations of hyperalignment and connectivity hyperalignment, which, for example, use iterative Procrustes transformations (Guntupalli et al., 2018, 2016; Haxby et al., 2011). We aim to describe the method in enough detail to provide a recipe for others. Our method was implemented using the Brain Imaging Analysis Kit (BrainIAK; https://brainiak.org), and code to perform these analyses is publicly available at https://github.com/snastase/connectivity-srm. To validate our approach, we first split each story in half; we designate the first half as the training set and the second half as the test set. All functional normalization algorithms are estimated from the training set and validated on the test set.

**Fig. 2.**
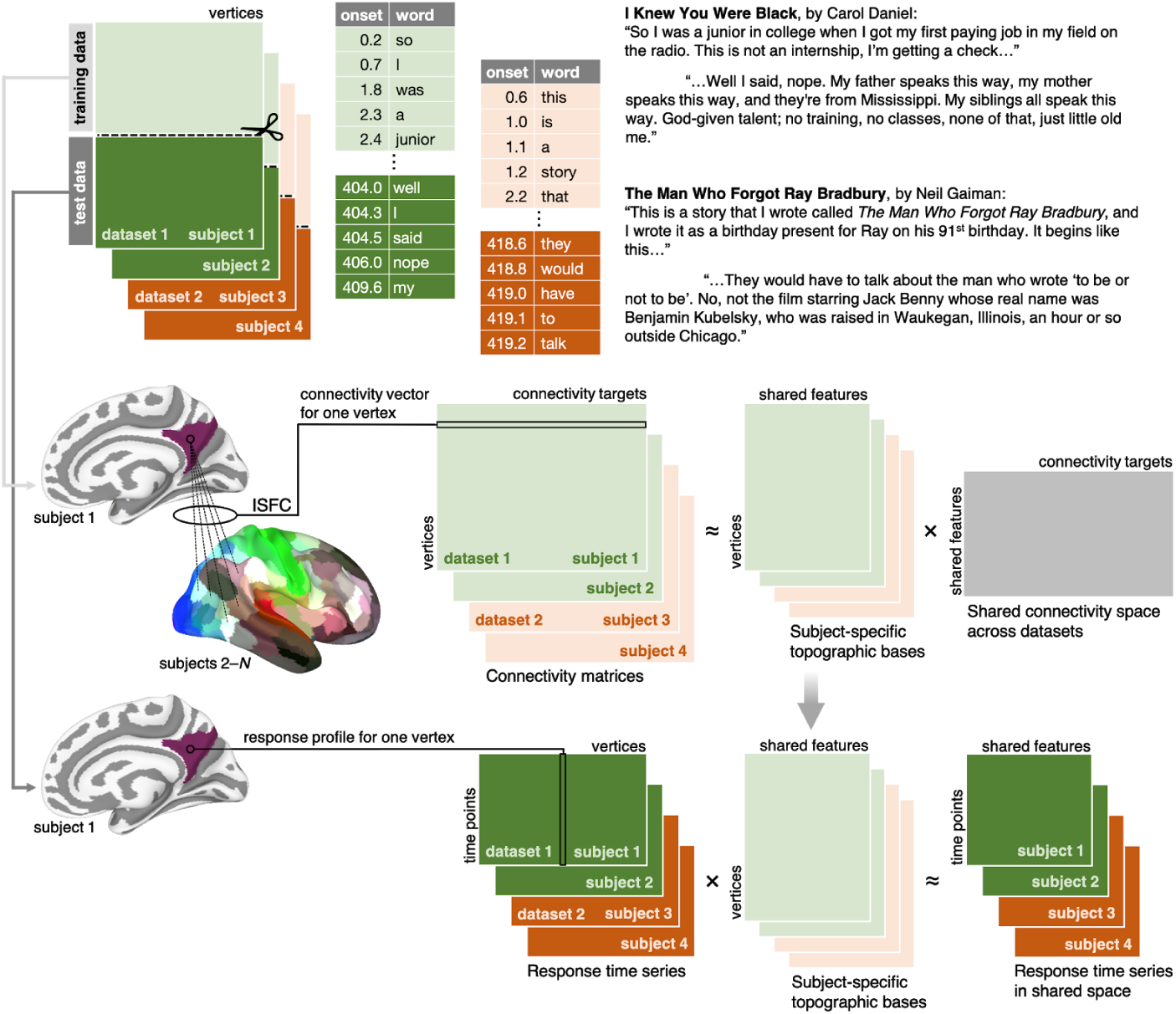
Schematic of connectivity-based shared response model. Data comprise multiple stories (e.g., dataset 1–2) and largely non-overlapping samples of subjects (e.g., subject 1–*N*). Time-stamped transcripts are shown for example stories “I Knew You Were Black” by Carol Daniel and “The Man Who Forgot Ray Bradbury” by Neil Gaiman. Data were partitioned into training and test sets for cross-validation: the first half of each story was assigned to the training set (light colors), and the second half was assigned to the test set (dark colors). For a given ROI, we computed ISFC between the response time series at each vertex and the average response time series for each of 360 cortical areas (connectivity targets) in the training set. We then used SRM to decompose these ISFC matrices into a set of subject-specific transformation matrices (topographic bases) and a single shared connectivity space across all datasets. We then apply the subject-specific transformations to response time series from the test set.

The following describes the cSRM algorithm for a given ROI. For each subject within a given dataset, we first estimate the functional connectivity between each voxel in the ROI and a set of connectivity targets. In the current context, because the subjects in a given dataset are all exposed to the same time-locked naturalistic story stimulus, we use ISFC to estimate stimulus-related functional connectivity and filter out idiosyncratic noise and intrinsic fluctuations (Nastase et al., 2019; Simony et al., 2016). Leave-one-subject-out ISFCs are computed by correlating the response time series in one subject with the average response time series across the remaining subjects in the same dataset (i.e., exposed to the same stimulus). To construct intersubject connectivity targets, we first extract for each subject the regional-average response time series for 360 areas spanning the cortex derived from a multimodal surface-based cortical parcellation (Glasser et al., 2016). For a given subject, we then average the 360 response time series across all other subjects (excluding the current subject). Finally, we compute the Pearson correlation between the response time series at each voxel in the ROI for the left-out subject and the regional-average response time series for 360 connectivity targets derived from the remaining subjects. This yields an asymmetric ISFC matrix for each subject with a number of columns corresponding to the number of voxels in the ROI and 360 rows representing the connectivity targets. The logic so far may seem counterintuitive; i.e., how can poorly-aligned connectivity targets be used to “bootstrap” fine-grained alignment? This approach hinges on the assumption that the voxels within an ROI are characterized by topographic “signatures” of long-range functional connectivity (Arcaro et al., 2015; e.g., Heinzle et al., 2011; Jbabdi et al., 2013). Connectivity targets are deliberately coarse (to mitigate deficiencies of anatomical alignment), but widespread enough to afford distinct connectivity signatures for driving fine-grained alignment. Although the overall aim is similar, our method of estimating connectivity differs from the core implementation of connectivity hyperalignment by Guntupalli and colleagues (2018) in several ways. First, we estimate functional connectivity using ISFC rather than within-subject functional connectivity. ISFC analysis relies on a shared stimulus to effectively isolate stimulus-related connectivity and is not applicable to resting-state data. Second, Guntupalli and colleagues (2018, pp. 20–21) apply connectivity hyperalignment within each connectivity target prior to computing connectivity vectors for the voxels of interest. Instead we simply use the average time-series per target, limiting ourselves to coarser-grained, lower-dimensional connectivity vectors. Third, we use a predefined multimodal cortical parcellation to delineate connectivity targets rather than regularly-spaced searchlights agnostic to areal borders.

In contrast to time-series hyperalignment, which relies on a shared stimulus to evoke time-locked response trajectories across subjects, moving to connectivity abstracts away from the time series. Time-series hyperalignment operates on response matrices where the number of columns corresponds to the number of voxels or vertices in the ROI and the number of rows corresponds to the number of time points in the experiment. Each column corresponds to the response time series for a single voxel and each row corresponds to the distributed response pattern for a given time point. Different stimuli result in different response trajectories that cannot effectively be aligned and may yield different shared spaces (although these spaces may converge for sufficiently rich stimuli). On the other hand, connectivity hyperalignment operates on connectivity matrices where the number of rows instead corresponds to the number of connectivity targets elsewhere in the brain. In this framework, each column corresponds to the connectivity vector (a coarse whole-cortex connectivity profile) for a given “seed” voxel in the ROI; each row in the matrix corresponds to a fine-grained, spatially distributed pattern of connectivities across voxels in the ROI relative to a given connectivity target. Critically, the shape of the connectivity matrices is dictated by the number of connectivity targets, not that number of time points in a stimulus. These connectivity matrices are isomorphic across stimuli and can be more readily aggregated than disparate response trajectories. The “second-order isomorphism” of connectivity matrices allows us to aggregate (e.g., stack) ISFC matrices across both subjects and story stimuli.

The goal of hyperalignment is then to leverage commonality of function to find a set of transformations (e.g., rotations) that map each subject’s idiosyncratic voxel response space into an abstract, shared response space. The SRM implementation used here frames this in terms of matrix factorization (Anderson et al., 2016; Chen et al., 2015), where the aggregate data matrix across subjects is decomposed into a reduced-dimension shared space and a set of subject-specific topographic transformation matrices. Applying this algorithm to connectivity matrices yields a shared connectivity space. Intuitively, it may seem that the resulting subject-specific transformations are only suitable for aligning connectivity data—but this is not the case (Guntupalli et al., 2018). Importantly, these subject-specific matrices are topographic transformations and can be used to project response time series into the shared space—assuming the connectivity matrices capture sufficient functional commonality to effectively align response trajectories. Concretely, we applied the SRM to the ISFC matrices derived from the training data (first half of each story), resulting in a shared connectivity space and subject-specific transformations. We then used these transformations to project response time series from the test set (second half of each story) into the shared space for evaluation. Note that the SRM algorithm can differ in performance relative to, e.g., the iterative Procrustes transformations used by Guntupalli and colleagues (2018; see, e.g., Chen et al., 2015 for comparisons among algorithms).

For subjects participating in multiple datasets, we computed their connectivity matrices separately per dataset, then averaged these prior to estimating the SRM. This ensures that each subject only submits one connectivity matrix to the SRM and only receives one transformation matrix into shared space. Note that this is not strictly necessary; multiple connectivity matrices could be submitted for a single subject participating in multiple datasets, yielding multiple dataset-specific transformations—but we did not explore this alternative here.

We can also exclude a subset of stories and subjects when estimating the shared space. First, we estimate a shared connectivity space based on the training data from a subset of stories. In the schematic depicted in Fig. 2, this corresponds to excluding, e.g., “dataset 1” (green) entirely when estimating the shared space; i.e., the shared space is estimated from datasets (e.g., “dataset 2”, orange) comprising stimuli and subjects not included in dataset 1. Subsequently, we compute connectivity matrices on the training data for a set of left-out subjects and stories not used to estimate the shared space. Given the preexisting shared connectivity space and a connectivity matrix for a given left-out subject, we can solve for that subject’s topographic transformation into the predefined shared space (as described in Chen et al., 2015). We can then use this transformation to project the left-out subject’s test data into shared space. That is, in Fig. 2, we use the training half of dataset 1 (light green) to define a transformation into the preexisting shared space, and project the test half of dataset 1 (dark green) into this independent shared space. Projecting left-out subjects into a predefined shared space in this manner does not alter the shared space (or the preexisting transformations for subjects used to estimate the shared space). Note that in the context of connectivity SRM, the training data used to project left-out subjects into shared space comprise connectivity matrices that are not strictly tied to a given stimulus; this allows novel subjects listening to novel stories to be transformed into an independent, predefined shared space.

Although we focus on defining a single shared connectivity space across datasets, we can also apply cSRM separately to each story dataset in isolation to create story-specific shared spaces. We directly compare the single connectivity-based shared space defined across all stories to story-specific connectivity-based shared spaces defined separately for each dataset. We also compare cSRM to conventional within-story time-series hyperalignment using the analogous SRM implementation (tSRM; Chen et al., 2015). SRM yields a reduced-dimension shared space at a specified dimensionality of *k* shared features; in several cases we compare shared spaces at varying dimensionality. To control for uninteresting effects of dimensionality reduction with SRM, we also performed principal components analysis (PCA) at matched dimensionality (as in, e.g., Chen et al., 2015). PCA imposes an orthogonality constraint analogous to SRM, but when applied to the aggregated subject data yields the same projection across subjects. Intuitively, PCA can be thought of as a control condition implementing similar dimensionality reduction, but without accounting for topographic idiosyncrasies across subjects. When interpreting results, cSRM performance should be compared to PCA at the matching dimensionality.

### Time-segment classification

We evaluated cSRM against alternative normalization schemes using between-subject story time-segment classification (Haxby et al., 2011). This analysis measures how accurately brief spatiotemporal response trajectories corresponding to unique segments of a story can be matched across subjects. We divided the test data (second half of each story) into 10-TR (15-second) segments and concatenated the response patterns across TRs into a single spatiotemporal response vector (or response trajectory) per segment. To perform between-subject classification for a given test subject, we first averaged the response vectors for each time segment over *N*–1 subjects excluding the test subject. We then computed the Pearson correlation between each response vector in the test subject and the average response vectors from the remaining subjects. A given response vector in the test subject was correctly classified if it is most highly correlated with the correct average vector from the remaining subjects. This is effectively a correlation-based 1-nearest neighbor classifier with leave-one-subject-out cross-validation (Haxby, 2012). Chance accuracy is 1 over the number of time segments in the test data for a given story and varies across stories. Note that, by design, this analysis can capitalize on any information shared across subjects and is agnostic to the type of information (e.g., sensory, semantic) encoded in response trajectories.

### Intersubject correlation

We used two varieties of intersubject correlation analysis to further dissect how cSRM affects spatially distributed response time series. We first examined how cSRM impacts vertex- or feature-wise intersubject time-series correlations (Hasson et al., 2010, 2004; Nastase et al., 2019). For each subject, we computed the Pearson correlation between the response time series at each vertex or feature and the average response time series at that vertex or feature across *N* – 1 remaining subjects (i.e., leave-one-out temporal ISCs). For each vertex or feature, we then averaged ISCs across subjects to summarize the shared signal for that vertex/feature. To evaluate how cSRM affects spatially distributed response patterns, we computed intersubject pattern correlations at each time point (Chen et al., 2017; Nastase et al., 2019; Zadbood et al., 2017). More specifically, for each subject, we computed the Pearson correlation between the response pattern at each TR and the average response pattern for *N* – 1 remaining subjects, then averaged these correlations across all time points per subject (i.e., leave-one-out spatial ISCs). We summarized these values by averaging the resulting ISCs across time points. Note that cSRM could conceivably produce the same degenerate response pattern across all time points and still yield high spatial ISCs, despite effectively discarding the stimulus-specific information of interest. However, this would be inconsistent with high time-segment classification accuracies; therefore, we consider spatial ISCs as a view into the observed time-segment classification performance rather than a benchmark in isolation.

### Semantic encoding model

We also evaluated how cSRM impacts model-based encoding and decoding for two stories (Fig. 3; Güçlü and van Gerven, 2017; Van Uden et al., 2018; Vodrahalli et al., 2017; Wen et al., 2018). To quantify the semantic content of the stories, we first used a semi-supervised forced-alignment algorithm (Yuan and Liberman, 2008) to extract time-stamped transcripts from each story stimulus (see Fig. 1 for an example). We then assigned semantic vectors to each word from the 300-dimensional word2vec embedding space trained on ∼100 billion words from the Google News corpus (Mikolov et al., 2013). More semantically similar words are located nearer to each other in this vector space; that is, they have more similar word embeddings (Turney et al., 2010). For each TR, all words with onsets occurring within that TR were assigned to the TR, and any words spanning two TRs were assigned to both. For TRs containing multiple words, we simply averaged the corresponding word embeddings to produce a single semantic vector per TR (cf. Vodrahalli et al., 2017). TRs in which no words occurred were assigned zero vectors. To account for varying hemodynamic lag, the 300-dimensional model was concatenated at delays of 2, 3, 4, and 5 TRs (3.0, 4.5, 6.0, 7.5 s), yielding a 1200-dimensional vector per TR (similarly to Huth et al., 2016). Delays were applied separately for the training and test data so as to avoid leakage across the train–test boundary.

**Fig. 3.**
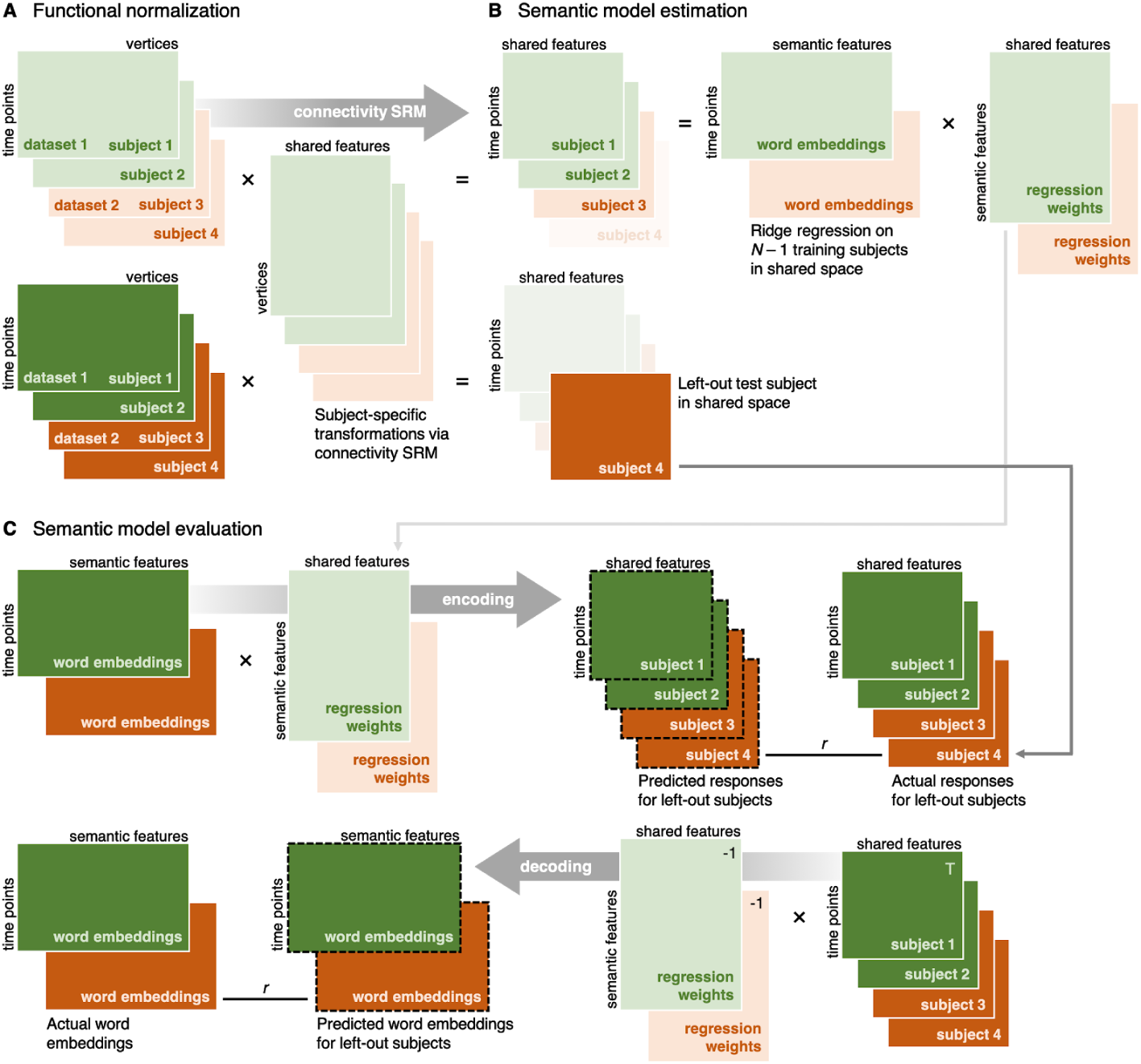
Schematic of semantic model-based encoding and decoding. Connectivity SRM is used to project both the training and test data into the shared space. Light colors indicate training data (first half of each story) and dark colors indicate test data (second half of each story; as in Fig. 2). Here we average response time series for the *N* – 1 training subjects. Ridge regression is then used to find a set of coefficients (weights) mapping from the semantic feature space (word embeddings) to the response time series at each vertex or feature. In the forward encoding analysis, we use the weight matrix estimated from the training data to predict vertex- or feature-wise response time series from the semantic vectors in the test set. In the decoding (or inverse encoding) analysis, we use the inverse of this weight matrix to predict semantic vectors from the response patterns at each time point. In both cases, we use correlation to evaluate that match between the predicted and actual response time series or semantic vectors.

To assess whether semantic information encoded in the embedding space captures variability in brain activity, we used a forward encoding model (Huth et al., 2016; Mitchell et al., 2008; Pereira et al., 2018; Wehbe et al., 2014). Following work by Huth and colleagues (2016), we used ridge regression to estimate coefficients (weights) for these 1200 semantic model features. *L*2-regularized linear regression effectively imposes a prior on the feature weights: the identity matrix scaled by a ridge coefficient (Diedrichsen and Kriegeskorte, 2017). Ideally, the optimal ridge coefficient is selected using nested cross-validation. However, this is computationally intensive, and will tend to yield different ridge coefficients for each subject, voxel, and cross-validation fold, complicating model comparison. For example, Huth and colleagues (2016) used a resampling approach to find optimal ridge coefficients, then averaged these across voxels and subjects to arrive at a single, consensus ridge coefficient (183.3 in that case). Here, to simplify numerous model comparisons, we use an arbitrary ridge coefficient of 100 throughout. This of course handicaps the absolute performance of our model. Our goal is not to engineer a novel or high-performing encoding model (see, e.g., Huth et al., 2016; Pereira et al., 2018), but to explore how functional normalization algorithms such as cSRM impact model performance under simplifying assumptions. We are interested in the encoding model insofar as it can provide insights into the performance of normalization algorithms.

Ridge regression was used to estimate weights so as to best predict the response time series at each vertex (or feature) in the training data (implemented using scikit-learn; Pedregosa et al., 2011). In the case of functional normalization (e.g., cSRM), note that we first estimated transformations into shared space from the training data. In order to fit the semantic encoding model, we used these transformations to project the training data (from which the transformations were derived) into shared space. That is, both the SRM transformations and the semantic model weights were estimated from the training data (first half of each story), affording unbiased validation on the test data (second half of each story). We focus on leave-one-subject-out cross-validation to evaluate the semantic encoding models: for each cross-validation fold, regression weights were estimated on the training data for *N* – 1 subjects, and evaluated on the test data for the left-out subject. In the current work, we average the training data across the *N* – 1 subjects in shared space before training; however, these data could be concatenated instead.

The regression weights estimated from the training data can then be used to predict response time series from the semantic vectors in a left-out subject. This approach—predicting vertex-wise response time series from the semantic model—is referred to as “forward encoding”. To evaluate the quality of these predictions, we computed the Pearson correlation between the predicted response time series and the actual response time series. We also perform a model-based decoding analysis, referred to as an “inverted encoding model,” to predict semantic vectors from response patterns in the test set (Brouwer and Heeger, 2009; Gardner and Liu, 2019; Sprague et al., 2018; Thirion et al., 2006). Note that this approach differs from generic classification analyses (Naselaris and Kay, 2015; e.g., Norman et al., 2006), which do not specify an explicit feature space for decoding. We first averaged coefficients across the four delays to obtain a single 300-dimensional weight matrix (as in Huth et al., 2012). We then computed the pseudo-inverse of the weight matrix comprising all vertices or features in an ROI. We multiplied the response pattern at each time point by this inverted weight matrix, resulting in a predicted semantic vector for each time point. We evaluated the quality of these predictions by computing the Pearson correlation between the predicted semantic vector for each time point and the actual semantic vectors for all time points in the test set. We then assess the rank of the correct semantic vector in the test set and normalize this by the number of semantic vectors in the test set to obtain a normalized rank accuracy score (Pereira et al., 2018). We average these rank accuracies across all time points in the test set. The rank accuracy score ranges from 0–1 where a score of 1 indicates that the correct semantic vector was the most similar to the predicted semantic vector and thus the highest-ranked vector. If there were no systematic relationship between predicted and actual semantic vectors, this would yield a chance rank accuracy score of approximately 0.5.

## Results

### Time-segment classification

We evaluated functional normalization algorithms in terms of between-subject story time-segment classification (Haxby et al., 2011). We divided the test data into 10-TR (15-second) response trajectories and supplied these to a between-subject correlation-based classifier (chance is 1 over the number of time segments for a given story). We first assessed between-subject time-segment classification in AAC across all 10 story stimuli (see Fig. 4). We compared classification performance for several implementations of functional normalization against the anatomically normalized data (“no SRM”), including time-series SRM defined within each story, connectivity SRM defined within each story, and connectivity SRM defined across all stories. For this representative example, we fit the SRMs with *k* = 100 shared features. We estimated 95% bootstrap confidence intervals surrounding the mean classification accuracy across left-out subjects and hemispheres by resampling subjects with replacement. We avoid performing gratuitous null-hypothesis statistical tests, but note that, considered in isolation, cases in which the 95% confidence interval for the mean of one condition does not cross the mean of another imply statistically significant differences (at p < .05; Nakagawa and Cuthill, 2007). The visual depiction of confidence intervals does not account for the within-subjects design within a story, and is therefore more conservative than a paired test (Loftus and Masson, 1994).

**Fig. 4.**
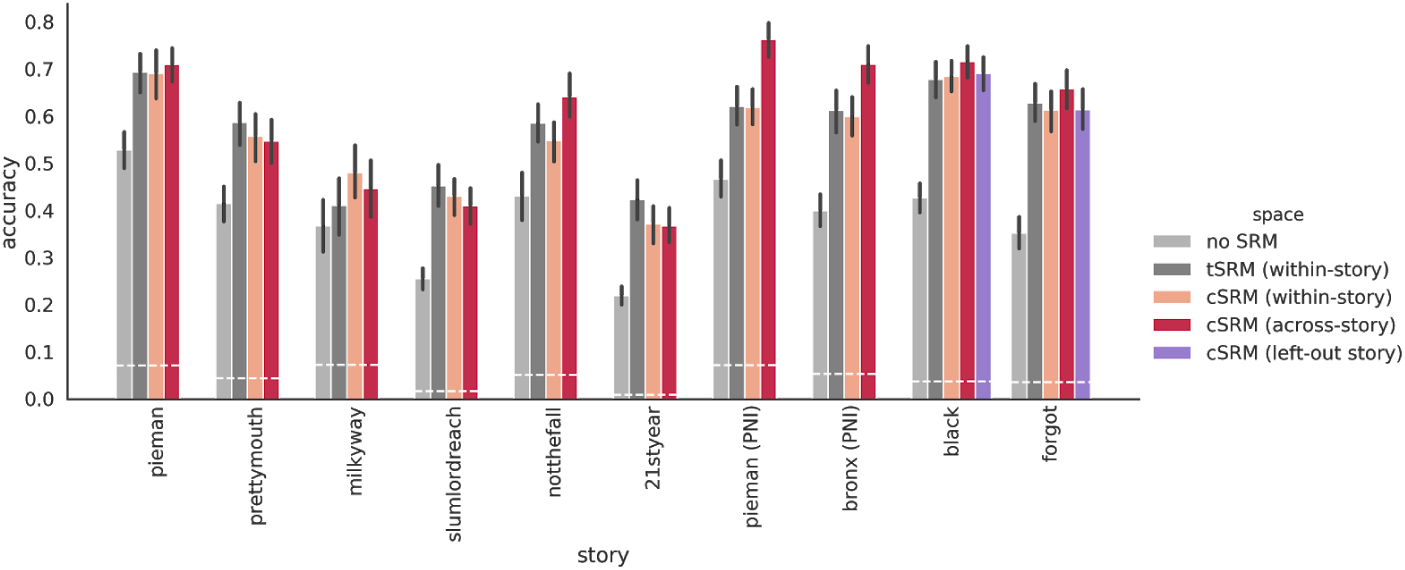
Time-segment classification across all stories in auditory association cortex (AAC). For each story, we compared surface-based anatomical alignment with no SRM (light gray), time-series SRM (necessarily defined within each story; dark gray), connectivity SRM defined separately within each story (pink), and a single connectivity SRM defined across all stories (red). We also recomputed a single connectivity SRM across all stories excluding subjects in the rightmost four datasets, and projected the “black” and “forgot” stories into this independent shared space (purple). The *y*-axis indicates between-subject time-segment classification accuracy averaged across left-out subjects and hemispheres. Dotted horizontal white lines indicate chance accuracy for each story (1 over the number of time segments in the test data). Error bars indicate 95% bootstrap confidence intervals estimated by resampling left-out subjects with replacement.

In general, all functional normalization algorithms provided considerable gains in time-segment classification over surface-based anatomical normalization. In most cases, the connectivity SRM was comparable to time-series SRM. Defining a connectivity-based shared space across all stories yielded comparable results to cSRMs defined separately for each story. In some cases—for example the “Pie Man” and “Running from the Bronx” stimuli recorded at PNI—the cSRM defined across stories provided marked improvement over other functional normalization algorithms. In general, classification performance on average improved from 40.3% with anatomical alignment to 63.7% with cSRM defined across all stories; tSRM and within-story cSRM yielded summary accuracies of 59.8% and 58.1% respectively.

Finally, we estimated a separate connectivity space across all stories excluding the four most recently collected stories (and the constituent subjects). We then computed connectivity matrices from the training data for each left-out subject in the left-out stories “I Knew You Were Black” and “The Man Who Forgot Ray Bradbury”. Given the predefined shared space and a connectivity matrix for each subject, we can derive a transformation for each left-out subject into the preexisting shared space (Chen et al., 2015). Projecting the test data for left-out subjects into this independent shared space yielded comparable improvements in accuracy over anatomical alignment (from 39.0% with anatomical alignment to 65.3% with the independent cSRM). This suggests that (*a*) the shared space generalizes to novel subjects and stimuli, and (*b*) the connectivity estimates for the training half of these left-out stories are sufficient for aligning these data to the shared space. In addition to comprising a completely non-overlapping sample of subjects viewing a different stimulus, these data were collected on a different scanner model using a different acquisition sequence.

We next assessed between time-segment classification for two example stories (“I Knew You Were Black” and “The Man Who Forgot Ray Bradbury”) using cSRM at varying dimensionality across all four ROIs (Fig. 5). We compared classification performance for cSRM at dimensionalities *k* = 100, 50, and 10 shared features to anatomical normalization and PCA at matching dimensionality. PCA provides a control for the dimensionality reduction of SRM without resolving functional topographies across subjects. In EAC, cSRM afforded minimal improvements over anatomical alignment and only at low dimensionalities (from 35.3% to 42.5% at the best-performing *k* = 10 across both stories; chance ≈3.7%). However, in AAC and TPOJ, cSRM markedly improved classification performance over anatomical alignment: from 39.0% to 70.2% at the best-performing *k* = 50 in AAC; and from 22.9% to 48.5% at the best-performing *k* = 100 in TPOJ. Classification performance was only modestly improved in PMC (from 16.1% to 21.9% at *k* = 50). The PCA control analysis indicates that decreasing dimensionality biases time-segment classification performance upward, but that reduced dimensionality alone cannot account for the improvement due to cSRM. Furthermore, the consistent pattern of increasing performance with decreasing dimensionality for PCA suggests—particularly for AAC and TPOJ, which do not perform best at lowest dimensionality—that a higher-dimensional shared space (e.g., *k* = 100, 50) may encode information lost at lower dimensionality (e.g., *k* = 10) for some ROIs.

**Fig. 5.**
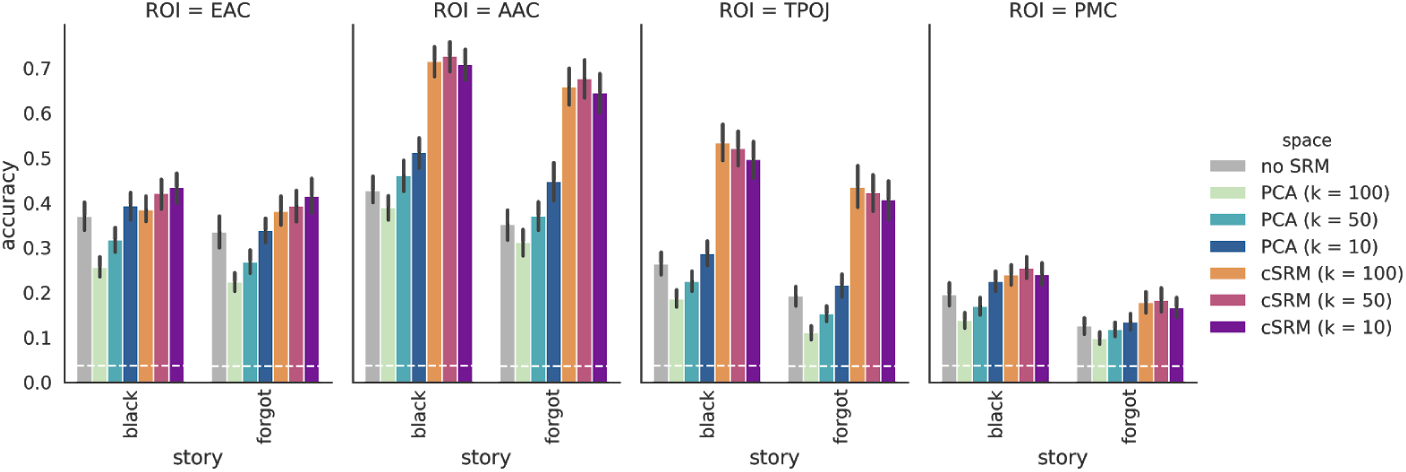
Time-segment classification at varying dimensionality for each ROI. For two example stories, we compared surface-based anatomical alignment with no SRM (gray), PCA controlling for the dimensionality reduction of SRM without resolving topographic idiosyncrasies (green–blue), and cSRM at dimensionalities *k* = 100, 50, and 10 (orange–purple). When interpreting reduced-dimension model performance, cSRM performance should be compared to PCA at the matching dimensionality. The *y*-axis indicates between-subject time-segment classification accuracy averaged across left-out subjects and hemispheres. Dotted horizontal white lines indicate chance accuracy for each story (1 over the number of time segments in the test data). Error bars indicate 95% bootstrap confidence intervals estimated by resampling left-out subjects with replacement.

### Intersubject correlation

The connectivity SRM maps each subject’s idiosyncratic responses into a shared space maximizing intersubject alignment of connectivity matrices—it does not explicitly optimize intersubject response time-series or pattern correlations. However, improvements in between-subject time-segment classification suggest that connectivity-based normalization in fact does implicitly align temporally-specific responses. To better understand this effect, we first examined how cSRM impacts response time series. For each subject, we computed vertex- or feature-wise intersubject time-series correlations (Hasson et al., 2004; Nastase et al., 2019). We compared temporal ISCs at each vertex with anatomical alignment alone, with PCA to control for dimensionality reduction in SRM, and with cSRM at dimensionalities *k* = 100, 50, and 10. This comparison is not straightforward: the anatomically-aligned ROIs contain many hundreds of correlated features; on the other hand, cSRM (or PCA) reduces this feature space to fewer dimensions accounting for orthogonal (uncorrelated) components of response variance (each feature reflects a distributed response topography across the entire ROI).

We first visualized the average ISC across subjects for every vertex or feature in the ROI (Fig. 6a). This reveals, for example, that cSRM and PCA isolate very few features in EAC (∼1 in each hemisphere) that capture the vast majority of the shared signal across subjects. In downstream ROIs, however, cSRM yields a considerably larger number of orthogonal dimensions capturing shared signal than PCA. This implies that functional alignment reveals higher-dimensional shared information across subjects. To more intuitively visualize this, we computed the number of features in each ROI exceeding an arbitrary ISC threshold of *r* > .1 (Fig. 6B). Note that in the reduced-dimension spaces, the absolute number of features exceeding this threshold is limited by the specified number of features *k*. We considered visualizing instead the proportion of features exceeding threshold relative to the maximum possible number of features *k*, but this obscures the fact that in most cases cSRM at, e.g., *k* = 100 yields several times the absolute number features exceeding threshold as cSRM at *k* = 10. These features represent orthogonal components of the response and a greater absolute number of features reflects higher-dimensional information shared across subjects. In AAC, for example, cSRM at *k* = 100 yields on average 79 features with ISC exceeding .1, while PCA at *k* = 100 yields only 21. Similarly, in TPOJ, cSRM at *k* = 100 increases the number of orthogonal features exceeding this threshold from 15 to 66. In PMC, cSRM at *k* = 100 increases the number of features exceeding threshold from 12 to 50. Even in EAC, cSRM at *k* = 100 yields over twice the number of features with ISCs exceeding .1 as PCA at matched dimensionality (39 and 16 features, respectively). Again, unlike vertices in anatomical space, these features represent distributed topographies across the ROI and capture orthogonal components of response variance.

**Fig. 6.**
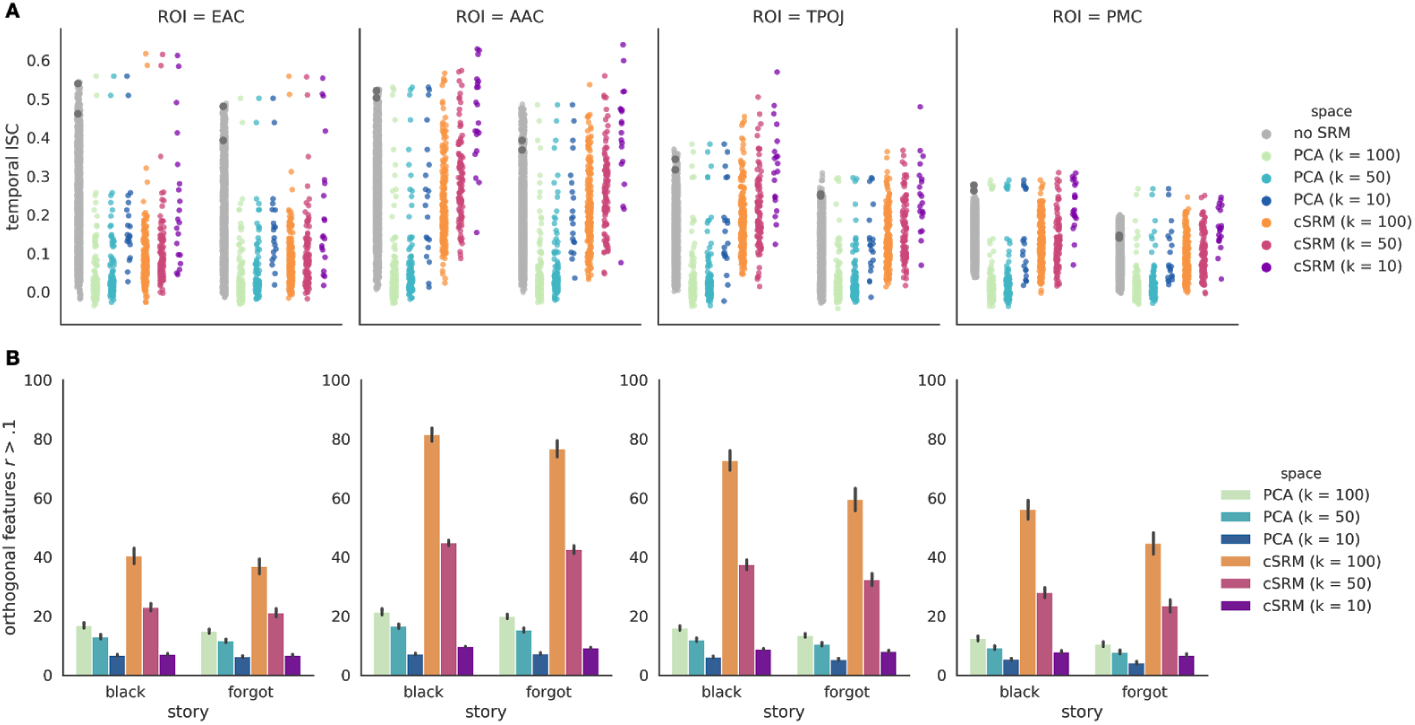
Intersubject time-series correlations per vertex/feature. (*A*) We computed the average ISC across subjects for each vertex with anatomical alignment and no SRM (light gray). Each marker corresponds to a vertex or feature for each hemisphere. We also computed ISCs on the regional-average response time series per hemisphere for the purpose of comparison (dark gray markers overlaid on the “no SRM” strip). We also visualize the average ISC across subjects for each feature after dimensionality reduction using PCA (green–blue), and cSRM at dimensionalities *k* = 100, 50, and 10 (orange–purple). When interpreting reduced-dimension model performance, cSRM performance should be compared to PCA at the matching dimensionality. The *y*-axis indicates the average temporal ISC across subjects per vertex or feature in each hemisphere. (*B*) We computed the number of orthogonal features with leave-one-out ISCs exceeding a threshold of *r* > .1 per subject (the maximum of which is limited by the specified *k*). The *y*-axis indicates the average number of features with ISCs exceeding this threshold across subjects. Note that the absolute number of features exceeding threshold is limited by the total number of features *k*. Error bars indicate 95% bootstrap confidence intervals estimated by resampling subjects with replacement.

Can the transformations derived from connectivity SRM align punctate, spatially distributed response patterns across subjects? To provide another window into the improvement in time-segment classification, we computed intersubject pattern correlations at each time point (Chen et al., 2017; Nastase et al., 2019; Zadbood et al., 2017). We assessed these spatial ISCs with anatomical alignment, PCA, and cSRM (Fig. 7). We found that spatial ISCs generally increased with reduced dimensionality, but that cSRM provides a boost over both anatomical alignment and PCA. In EAC, the benefit of cSRM over anatomical alignment was negligible, but exceeded PCA at matching dimensionality. In AAC and TPOJ, however, cSRM significantly improved the alignment of spatial response topographies across subjects. For example, in AAC, spatial ISCs almost, from *r* = .110 with anatomical alignment (*r* = .058 with PCA) to *r* = .211 with cSRM at *k* = 100. An effect of similar relative magnitude was observed in TPOJ: from *r* = .082 with anatomical alignment (*r* = .029 with PCA) to *r* = .153 at cSRM at *k* = 100. In PMC, spatial ISCs increased from *r* = .074 (*r* = .021 with PCA) to *r* = .100 with cSRM at *k* = 100.

**Fig. 7.**
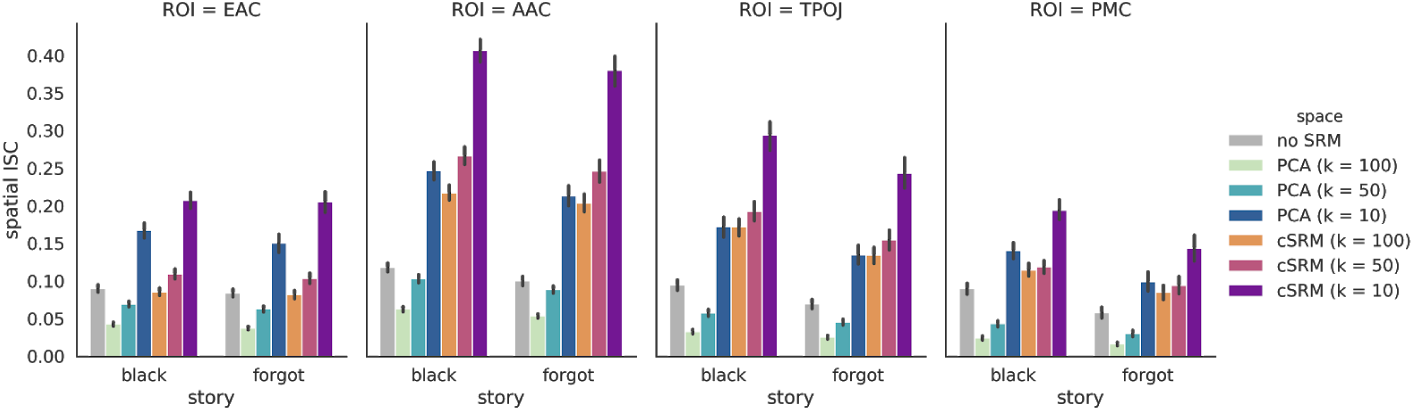
Intersubject pattern correlations. We compared spatial ISCs with anatomical alignment (gray), PCA to control for the reduced dimensionality of SRM (green–blue), and cSRM with dimensionalities *k* = 100, 50, 10 (orange–purple). When interpreting reduced-dimension model performance, cSRM performance should be compared to PCA at the matching dimensionality. The *y*-axis represents spatial ISC for each time point averaged across time points and subjects. Error bars indicate 95% bootstrap confidence intervals estimated by resampling subjects with replacement.

### Semantic encoding model

Encoding models have been increasingly adopted as a means of testing explicit feature spaces capturing representational content ranging from low-level visual features to high-level semantic content (Naselaris et al., 2011; Serences and Saproo, 2012). However, these models are often estimated independently per subject using large volumes of data (e.g., Huth et al., 2016), which poses problems of both scalability (in terms of data collection) and generalizability across subjects (cf. Güçlü and van Gerven, 2017; Van Uden et al., 2018; Vodrahalli et al., 2017). Here we used a simplistic semantic encoding model to explore how cSRM impacts model performance. We assigned semantic word embeddings to each time point, then used ridge regression to estimate a weight matrix mapping between word embeddings and brain responses (see Fig. 3).

In the forward encoding analysis, we used the regression weights estimated from the training data to predict vertex- or feature-wise response time series in the test set for a left-out subject. To evaluate the forward model, we computed the Pearson correlation between the predicted time series and actual time series for each vertex or feature. We compared forward encoding performance with surface-based anatomical alignment, PCA to control for dimensionality reduction in SRM, and cSRM at dimensionalities *k* = 100, 50, and 10. Although we emphasize a predictive approach using leave-one-subject-out cross-validation, we also present the typical within-subject performance (as in, e.g., Huth et al., 2016). Similarly to the temporal ISC analysis, this makes for a difficult comparison because vertices in the anatomical ROI are highly redundant (correlated), while PCA and cSRM reduce this feature space to fewer, orthogonal dimensions. We first visualized the vertex- and feature-wise performance averaged across subjects (Fig. 8A), then plotted the number of features in each ROI exceeding an arbitrary performance threshold of *r* > .1 (Fig. 8B). Although the maximum number of features exceeding threshold is limited by the specified *k*, greater absolute numbers of orthogonal features surpassing this threshold reflect higher-dimensional semantic information shared across subjects. In all cases, cSRM dramatically increased the number of well-predicted orthogonal features relative to the dimensionality-matched PCA control. In EAC, cSRM at *k* = 100 doubled the number of orthogonal features with performance exceeding *r* > .1 from 11 to 22. In AAC and TPOJ, cSRM at *k* = 100 roughly tripled the number of orthogonal features exceeding the forward encoding performance threshold—from 16 to 49 and from 12 to 36, respectively. In PMC, the number of features exceeding this threshold increased from 9 with PCA to 23 with cSRM at *k* = 100.

**Fig. 8.**
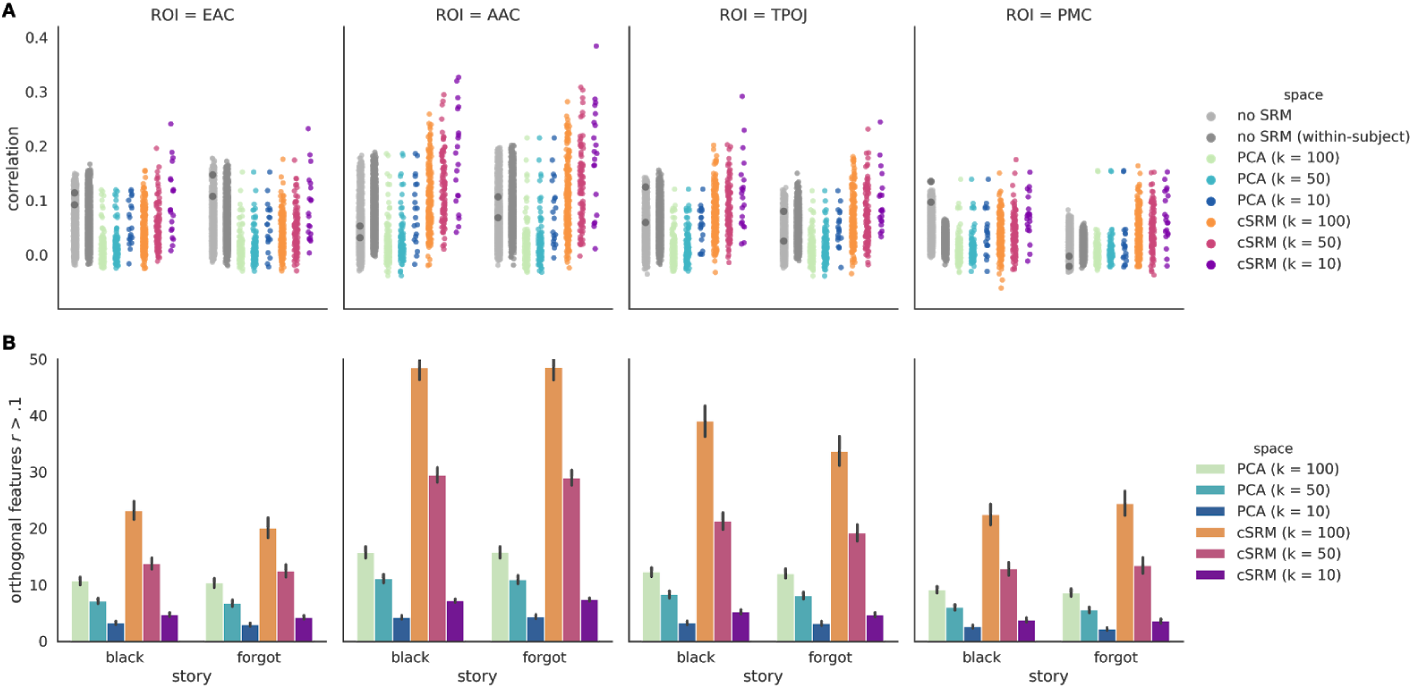
Forward encoding model performance. (*A*) We evaluated the vertex-wise between-subject (light gray) and within-subject (dark gray) forward encoding models with anatomical alignment (no SRM). Forward encoding performance for the regional-average response time series per hemisphere is visualized for comparison (dark gray markers overlaid on the “no SRM” strip). We also compared between-subject forward encoding performance for each feature after dimensionality reduction using PCA (green–blue), and cSRM at dimensionalities *k* = 100, 50, and 10 (orange–purple). When interpreting reduced-dimension model performance, cSRM performance should be compared to PCA at the matching dimensionality. The *y*-axis represents the average correlation between predicted and actual response time series across test subjects for each vertex or feature in each hemisphere. (*B*) We computed the number of orthogonal features with forward encoding performance exceeding a threshold of *r* > .1 per subject. The *y*-axis represents the average number of features performance exceeding this threshold across test subjects (and hemispheres). Error bars indicate 95% bootstrap confidence intervals estimated by resampling test subjects with replacement.

We next inverted the encoding model to predict semantic vectors from spatially distributed response patterns (Brouwer and Heeger, 2009; Gardner and Liu, 2019; Sprague et al., 2018; Thirion et al., 2006). While between-subject time-segment classification can capitalize on any diagnostic information shared across subjects, between-subject model-based decoding is strictly limited to the representational content encoded in the model. For each time point in the test set, we multiplied the distributed response pattern by the inverted weight matrix to recover a predicted semantic vector for that time point. To evaluate decoding performance, we computed the Pearson correlation between the predicted semantic vector and the actual semantic vectors in the test set and summarized this using a normalized rank accuracy score (Pereira et al., 2018). We compared between-subject model-based decoding using anatomical alignment, PCA, and cSRM (Fig. 9).

**Fig. 9.**
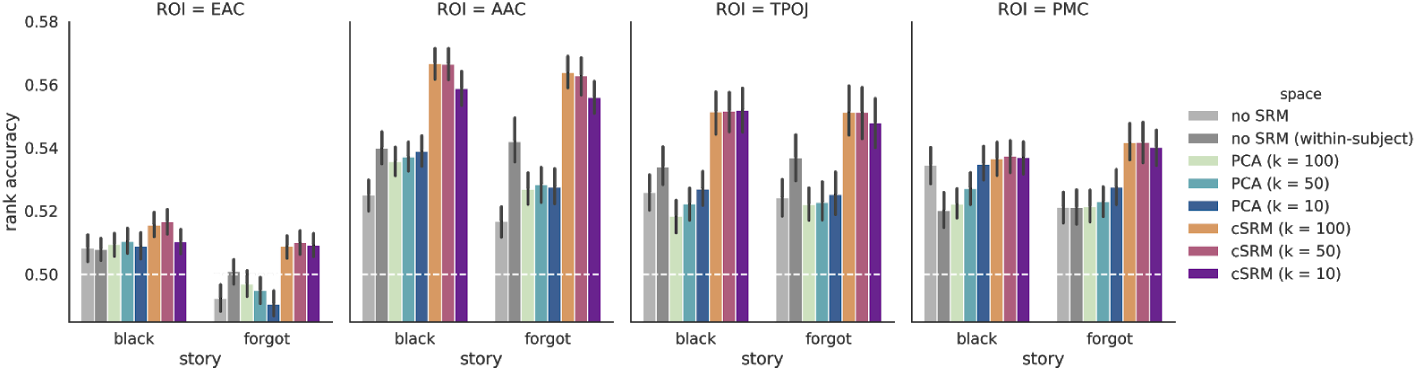
Model-based decoding performance. We compared between-subject decoding performance using anatomical alignment (light gray), PCA matching the dimensionality reduction of SRM (blue–green), and cSRM at dimensionalities *k* = 100, 50, and 10 (orange–purple). When interpreting reduced-dimension model performance, cSRM performance should be compared to PCA at the matching dimensionality. We also provide within-subject decoding performance for comparison (dark gray). The *y*-axis indicates rank accuracy averaged across time points, subjects, and hemispheres. Dotted horizontal lines indicate chance accuracy of approximately 50% for the rank accuracy score. Error bars indicate 95% bootstrap confidence intervals estimated by resampling test subjects with replacement.

Although we focus on between-subject decoding using leave-one-subject-out cross-validation, we also computed within-subject decoding performance for comparison. In general, model-based decoding accuracies were low; likely due to our simplistic model estimation procedure. However, we observed some interesting trends. First, within-subject performance exceeded between-subject performance using surface-based alignment in AAC and TPOJ, suggesting that anatomical normalization alone fails to translate some semantic information across brains. Second, dimensionality reduction did not consistently increase performance. Crucially, in almost all cases, between-subject decoding performance using cSRM exceeded both between- and within-subject performance using anatomical alignment. For example, in AAC, the average rank accuracy across test subjects increased from 52.1% with anatomical alignment (54.1% for within-subject decoding) to 56.5% with cSRM at *k* = 100 (with a rank accuracy of 63.5% for the best-performing test subject).

### Consensus space across stimuli

Finally, we examined the consequences of deriving a single shared space across numerous distinct stories. Is the single, connectivity-based shared space defined across all datasets notably different from shared connectivity spaces derived separate for each story? For example, we may expect that each story considered in isolation will nonetheless yield similar shared spaces due to the nature of functional connectivity. To address this, we computed ISFC matrices from the test data for each subject and projected these ISFC matrices into either (*a*) the connectivity-based shared space defined across all stories (across-story cSRM), or (*b*) the unique connectivity-based shared spaces defined separately for each story (within-story cSRM). For illustrative purposes, we used AAC and cSRM at *k* = 100. We then flattened and averaged the ISFC matrices across subjects per story in their respective shared spaces and computed the pairwise correlations between the mean ISFC matrices across stories (Fig. 10). In fact, ISFCs projected into the single, shared connectivity space are more similar across all pairs of stories than those projected into story-specific shared spaces: the average correlation of ISFC matrices increased from *r* = .596 to *r* = .756 across all pairs of stories. This is to be expected but serves as a useful sanity check, and suggests that the transformations defined across all stories point to a consensus space.

**Fig. 10.**
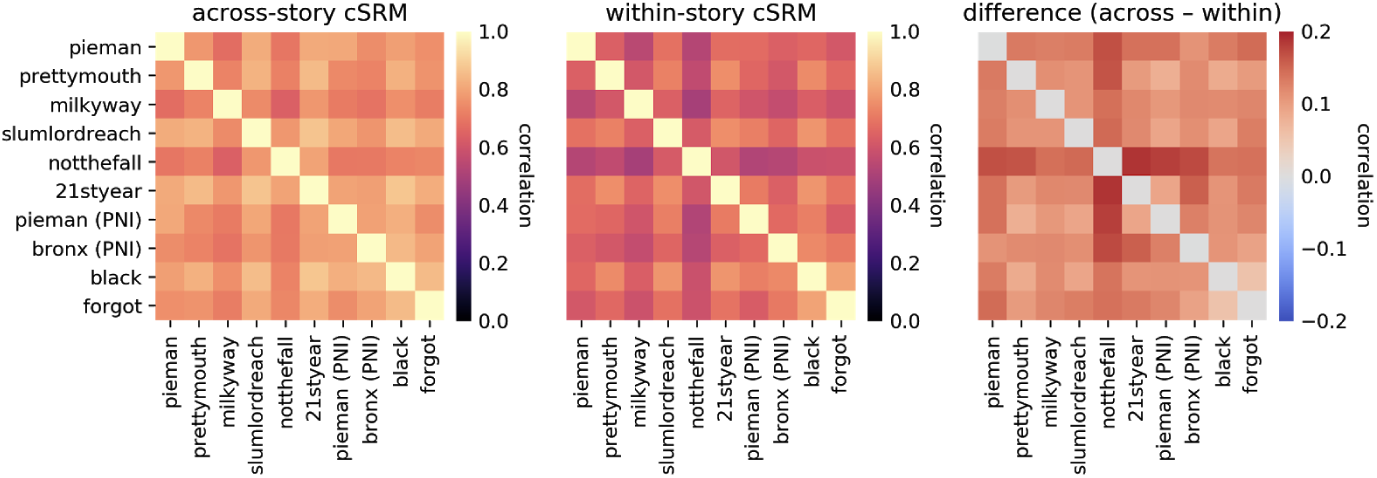
Story similarity in consensus and story-specific shared spaces. We projected ISFCs estimated from the test data into either a shared connectivity space defined across all stories (across-story cSRM) or story-specific shared connectivity spaces (within-story cSRM). We then computed the pairwise correlations of ISFC matrices across stories. The difference between these cSRMs (right) indicates that the cSRM defined across all stories projects into a consensus space.

We next considered whether constructing a shared space across distinct stimuli impacts responses within a story. Similarly to before, we projected the response time series for the test data into either a shared connectivity space defined across all stories or story-specific shared connectivity spaces (in AAC, using cSRM at *k* = 100). We then concatenated these response patterns over time into a single spatiotemporal response trajectory for each subject, and computed leave-one-subject-out ISCs on these spatiotemporal vectors within each story (Fig. 11; Nastase et al., 2019). Interestingly, we found that the response trajectories were more similar across subjects within a given story when projected into the shared space defined across stories. This increase in similarity was small, from *r* = .261 to *r* = .286, but statistically significant across all stories (*p* ≤ .001 for all stories, nonparametric Wilcoxon signed-rank test, Bonferroni corrected).

**Fig. 11.**
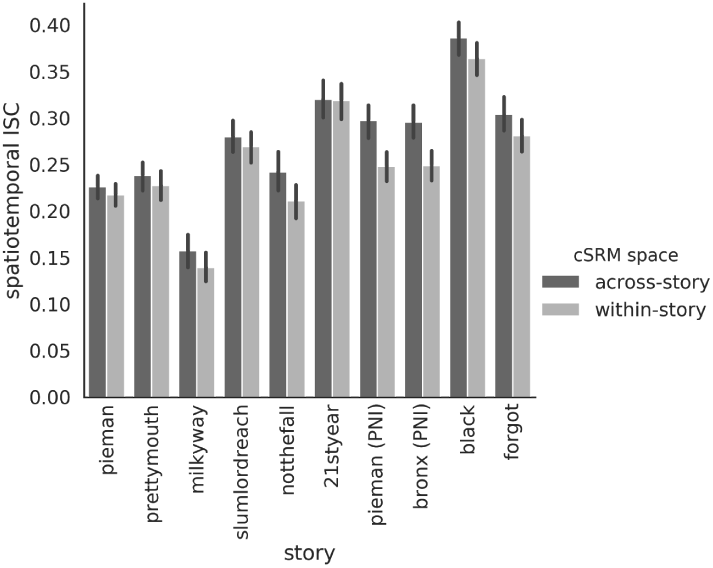
Intersubject spatiotemporal correlations in consensus and story-specific shared spaces. We projected spatiotemporal response trajectories into either a shared space defined across all stories (across-story) or story-specific shared spaces (within-story). We then computed the intersubject correlations of these response trajectories within each story. The *y*-axis indicates the average spatiotemporal ISC across subjects and hemispheres. Error bars indicate 95% bootstrap confidence intervals estimated by resampling subjects with replacement.

## Discussion

We have demonstrated that connectivity hyperalignment can be used to estimate a shared response space across disjoint datasets with unique stimuli and non-overlapping samples of subjects. Although the shared space is defined in terms of connectivity, the subject-specific topographic transformation matrices are suitable for projecting response time series into this space (Guntupalli et al., 2018). We have shown that transformations derived from intersubject functional connectivity are sufficient to precisely align response trajectories in a way that is both spatially and temporally specific; in the case of time-segment classification, this effect was consistent across all 10 stories. The features in the shared space derived using cSRM reflect orthogonal components of response variability; projecting data into shared space dramatically increases the dimensionality of information shared across subjects. The consensus space introduced here conjoins diverse story stimuli, and effectively regularizes subject- and story-specific transformations.

There are three key concepts underlying the success of this algorithm. First, relatively coarse connectivity targets spanning cortex provide can provide sufficiently rich signatures of functional connectivity to drive fine-grained topographic alignment (Arcaro et al., 2015; Heinzle et al., 2011; Jbabdi et al., 2013). Here we define connectivity targets according to a multimodal cortical parcellation (Glasser et al., 2016), which yields a relatively low-dimensional, more computational efficient connectivity space. However, there are a variety of ways to construct connectivity targets (cf. Guntupalli et al., 2018); for example, in the context of naturalistic stimuli, vertices with temporal ISCs exceeding some threshold could serve as potentially finer-grained connectivity targets. The second, related concept is that constructing a shared space based on functional connectivity yields subject-specific topographic transformations suitable for aligning response time series.

This may seem counterintuitive, but, again, suggests that long-range connectivity profiles capture information about local response topographies. Third, all datasets used here, despite showcasing story stimuli with diverse topics and speakers, are all examples of the superordinate story-listening “task.” This common task may yield commensurate connectivity patterns across stimuli, allowing the algorithm to find a consensus shared space. Future work is required to explore the boundary conditions of this algorithm and determine whether a shared space can be defined across qualitatively different tasks or paradigms.

Our analyses demonstrate that projecting data into a shared space derived from functional connectivity improves semantic model-based encoding and decoding. Interestingly, between-subject semantic model-based decoding with cSRM exceeded within-subject decoding in all ROIs. How can this be possible? It may be that our simplistic semantic encoding model may only capture relatively coarse-grained information across subjects. However, between-subject encoding models allow us to leverage much larger volumes of data than could be acquired in a single subject. Aggregating data across subjects can yield much cleaner training samples, thus improving performance over limited and often noisy within-subject data. Here, we fit the semantic encoding model on the averaged response time series across training subjects; although averaging yields clean response time series, subjects in the shared space could instead be concatenated, dramatically increasing the number of training samples available for modeling. Furthermore, our rudimentary fitting procedure (e.g., using an arbitrary, fixed ridge coefficient) will necessarily yield suboptimal model performance. Although we adopt this approach to reduce the computational burden and simplify model comparison, it could be argued that models are not fairly compared unless they can arrive at their own optimal ridge coefficients. For any use-case where evaluating or comparing encoding models is the primary scientific goal, we recommend grid search using nested cross-validation or resampling procedures to identify optimal hyperparameters (as in, e.g., Huth et al., 2016).

Interestingly, our functional normalization algorithm differentially benefited certain ROIs. For example, EAC, an early sensory ROI, was only marginally improved by cSRM, while downstream association cortices such as AAC and TPOJ were more dramatically improved. On the other hand, the putatively high-level PMC was only modestly improved by cSRM. There are several possible reasons for these discrepancies. First, the quality of the cSRM derives from the richness of functional connectivity for a given ROI; early sensory areas may have limited connectivity to the rest of the brain, whereas association cortices are in effect defined by their broad, integrative connectivity. Second, some cortical areas may have relatively stereotyped functional architecture across individuals or inherently coarse response topographies; both scenarios would reduce the benefits of functional normalization over anatomical normalization. Third, cortical areas vary in the extent to which their processing is strictly stimulus-locked; early sensory areas may be better aligned by temporal hyperalignment, while association cortices may be better aligned by connectivity hyperalignment (Guntupalli et al., 2018).

The approach described here has several limitations. We limit our analysis to a handful of ROIs and do not provide a whole-cortex searchlight-based solution as in the core implementation of connectivity hyperalignment (Guntupalli et al., 2018). However, it would be straightforward to extend the implementation used here to parcels tiling the entire cortex. Unlike Guntupalli and colleagues, we take advantage of the shared story stimulus within each dataset and use ISFC to filter out idiosyncratic, intrinsic fluctuations and isolate stimulus-related connectivity (Simony et al., 2016). This approach, however, is not applicable to resting-state paradigms where there is no shared stimulus. Furthermore, we do not currently account for the fact that certain stories have considerably more subjects than others. In our implementation of cSRM, this will tend to bias the shared space toward the stories with the most subjects. The contribution of each story to the shared space could conceivably be normalized by the proportion of subjects for that story relative to the total number of subjects. However, in practice we may want to bias the shared space toward stories with the largest number of subjects, as these may provide the most robust shared model for limited data.

This raises the important question of whether it is worthwhile to “sacrifice” functional data for the purposes of normalization. We generally advocate for estimating a shared space on independent data to avoid circularity (Kriegeskorte et al., 2009). In the current work we estimate the shared space from the first half of all stories and use the second half for evaluation, potentially undermining generalizability across stories. However, we demonstrate that left-out stories still benefit from connectivity SRM defined on independent data. On the other hand, many analyses (e.g., model-based encoding and decoding) already require independent training data for model estimation. Here we adopt an approach where both functional normalization and encoding model weights are estimated from the same training set, effectively negating the price paid for normalization.

Precision neuroscience is fundamentally limited by the feasibility of collecting large volumes of data in experimental subjects (let alone patients). The fact that connectivity hyperalignment benefits a completely independent set of subjects and stories not used to estimate the shared space has important implications. This capacity for generalization suggests that many existing datasets, from resting-state to naturalistic movies, could benefit from connectivity hyperalignment. This provides a means for using existing data to “bootstrap” improvements in functional registration and build better, more generalizable predictive models.

## Acknowledgments

We thank James V. Haxby, Feilong Ma, Eshin Jolly, Asieh Zadbood, Kristina M. Rapuano, and Erez Simony for helpful comments, and Janice Chen, Christopher Honey, Yaara Yeshurun, and Claire Chang for sharing previously-collected data.

